# Distinct Molecular Mechanisms Regulate Feeding State-Dependent CO_2_ Chemotaxis Plasticity During Different Life Stages in *Caenorhabditis elegans*

**DOI:** 10.64898/2026.02.06.704338

**Authors:** Akankshya Ramkrishna Sahu, Swarupa Mallick, Atal Vats, Abhishek Bhattacharya

**Affiliations:** National Centre for Biological Sciences - TIFR, Bangalore, India

## Abstract

Across developmental stages, animals modulate their behavioural responses to external cues according to intrinsic physiological states. During development, particularly in juvenile stages, nervous systems undergo extensive changes at multiple levels. However, it remains unclear whether nervous systems at different developmental stages employ the same underlying molecular mechanisms to produce equivalent behavioural modulations in response to intrinsic or extrinsic cues. Using the model organism *Caenorhabditis elegans*, we identify here that animals employ distinct molecular mechanisms to achieve equivalent modulation of CO_2_-chemosensory behaviour at different developmental stages. Ubiquitin-proteasome-mediated downregulation of the insulin/IGF receptor, DAF-2 by the conserved quality-control ubiquitin ligase CHN-1/CHIP promotes attraction to environmental CO_2_ during the starvation-induced L1-arrest stage. In contrast, CO_2_-attaction in dauer animals is independent of CHN-1/CHIP activity. Furthermore, the feeding-induced reversal of CO_2_-chemotaxis to avoidance during L1-arrest exit requires the insulin/IGF pathway and the conserved CRH-1/CREB1 transcription factor activity in the CO_2_-sensing BAG neurons. However, induction CO_2_-avoidance during dauer exit is independent of CRH-1/CREB1 activity. These findings suggest that neural circuits at different life stages may utilize distinct, stage-specific molecular mechanisms to induce identical plasticity in chemosensory behaviour.

## INTRODUCTION

Modulations of chemosensory behaviours in response to intrinsic physiological states or extrinsic environmental conditions are essential for successful navigation through complex environments and, ultimately, for survival across animal species. It has been shown that internal physiological states in both invertebrates and vertebrates can alter behavioural responses or brain activity to specific stimuli^1-6^. During development, particularly in juvenile stages, nervous systems undergo extensive changes at anatomical, gene expression, and functional levels, which have been associated with distinct behavioural responses to chemosensory cues. However, the molecular mechanisms underlying how nervous systems across developmental stages implement equivalent behavioural modulations in response to changes in intrinsic physiological states or extrinsic environment remain unclear.

Hunger plays a significant role in modulating animals’ response to external cues^1,7,8^. The insulin/IGFR-like signalling (IIS) pathway is a key determinant of feeding state-dependent modulation of metabolism, nervous system physiology, and behaviour in metazoans^9-11^. In fed animals, activation of the insulin/IGF-receptor upregulates the conserved phosphoinositide 3-kinase (PI3K) pathway, resulting in inhibition of the Forkhead transcription factor, DAF-16/FOXO. In contrast, in starved animals, downregulation of the IIS pathway leads to DAF-16/FOXO activation, which regulates many target genes in diverse tissue types to modulate various physiological processes^10,11^. The quality-control ubiquitin ligase CHIP has been shown to regulate the IIS pathway in nematode *C. elegans*, Drosophila, and humans, by directly targeting the insulin/IGF receptor^12^. Notably, the sole ortholog of CHIP in *C. elegans,* CHN-1, differentially regulates insulin/IGF receptor, DAF-2, at distinct developmental stages^12,13^.

The highly conserved transcription factor, cAMP response element binding protein (CREB1), also functions as a crucial sensor of internal physiological states and is involved in regulating various physiological processes across phyla, including metabolic control, energy homeostasis, aging, and memory formation ^14-18^. However, it remains to be fully understood whether CREB1 activities in the nervous system are modulated across different developmental stages.

The nematode *C. elegans* has a well-characterized nervous system and generates robust, well-defined behavioural responses to chemosensory cues. Moreover, the nervous system of *C. elegans* undergoes wide-scale anatomical and functional plasticity, resulting in markedly altered locomotory and chemosensory behaviours, when developing *C. elegans* larvae encounter acute environmental conditions during specific larval stages and develop into the dauer diapause stage^19-22^. Among the altered chemosensory preferences, the response to environmental CO_2_ concentration is completely reversed in dauer animals^22^. While well-fed adults exhibit strong avoidance towards CO_2_, dauers or adults under prolonged starvation display attraction to CO_2_^22-25^.

Apart from the dauer diapause stage, nutrient deprivation at the time of hatching or before molting into the L3 and L4 stages causes developmental arrest at specific checkpoint stages, namely, L1-, L3- and L4-arrest stages, respectively^26^. However, it remains unclear whether and how animals in these other developmentally arrested stages modulate CO_2_ chemosensory behaviour in response to the internal physiological state.

We describe here that distinct underlying molecular pathways can regulate identical plasticity of CO_2_ chemotaxis valence at different life stages. In addition to well-characterized dauers and starved adults, we identify that *C. elegans* in other developmental diapause stages also show CO_2_-attraction, which can be rapidly reversed to CO_2_ avoidance upon feeding. During other developmental arrest stages and in starved adults, the E3 ligase CHN-1/CHIP negatively regulates DAF-2 in a proteasome-dependent manner to promote CO_2_-attraction. In contrast, CHN-1 has no effect on CO_2_-attraction in dauers, which requires insulin/IGF-1 signalling activity^27^. Moreover, focusing on the L1-arrested animals, we show that both CO_2_-attraction and the attraction-promoting electrical synapse formation between BAG and AIB neurons are independent of DAF-16/FOXO transcription factors. Similarly, we show that distinct underlying pathways regulate feeding-dependent reversal of CO_2_-attraction in L1-arrested animals and in dauers. Although animals in both stages depend on insulin/IGF-1 signalling pathway activity for the onset of CO_2_-avoidance, L1-arrested animals specifically require the transcription factor CRH-1/CREB1 cell-autonomously in CO_2_-sensing BAG neurons. Taken together, our findings demonstrate that, depending on internal physiological states, neural circuits at distinct life stages utilize different molecular pathways to generate equivalent plasticity in chemosensory behaviour.

## RESULTS

### *C. elegans* across all developmental arrest stages exhibit CO_2_-attraction

Adult nematodes, including *C. elegans,* avoid CO₂ under well-fed conditions^22,23^. In contrast, long-term starved adults and dauer larvae of *C. elegans* display strong attraction to high concentrations of CO₂^22,23^. Among the dauer stage-specific alterations in the nervous system, the expression of a gap junction channel-forming innexin gene, *inx-6,* in the AIB interneuron leads to the formation of electrical synapses between AIB and the primary CO_2_-sensing neuron, BAG. This electrical synapse is shown to be required for optimal CO₂-attraction in dauers^19,28^ **(Figure 1A)**. However, in addition to the dauer stage, L1-arrested (L1d) animals also express *inx-6* in AIB neurons **(Figure 1A)**, leading to the formation of putative electrical synapses between BAG and AIB neurons^19^. These observations prompted us to test whether *C. elegans* at other developmental arrest stages also display altered CO_2_ chemotaxis valence.

**Figure 1:**
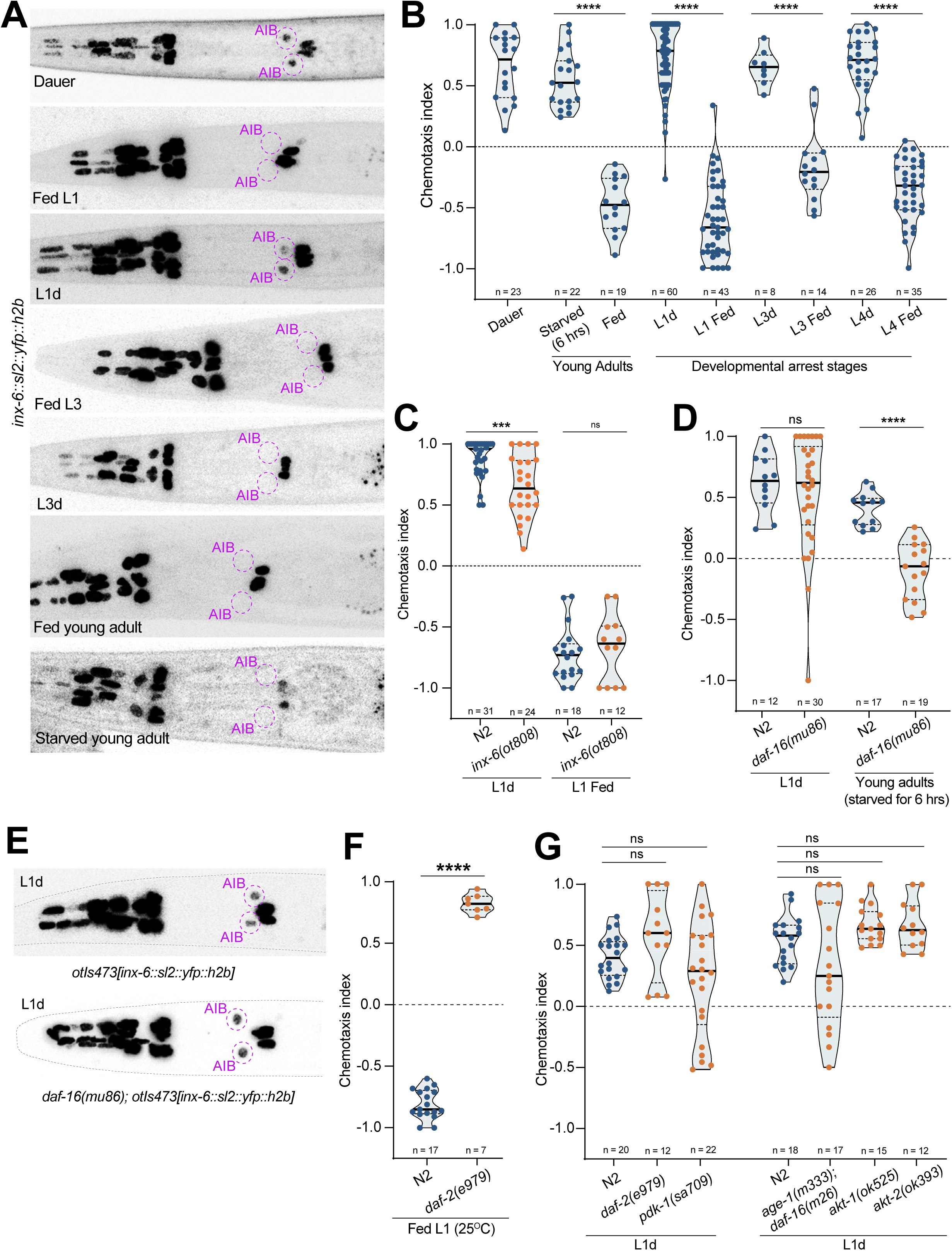
*C. elegans* in L1-, L3- and L4-arrest stages show CO_2_-attraction. (A) *inx-6* reporter allele, *inx-6(ot804)* is specifically turned on in AIB neurons in dauer and L1-arrest stages, but not during L3-arrest or in starved adult animals. (B-D and F-G) Each circle represents the chemotaxis index calculated from a single chemotaxis assay. Violin plots represent the distribution, black horizontal solid lines represent the median and horizontal dotted lines represent the quartiles. Mann–Whitney U tests p values for each comparison: n.s., nonsignificant, *p < 0.05, **p < 0.01, ***p < 0.001, ****p < 0.0001. (B) Similar to dauer and starved adult animals, L1-, L3- and L4-arrested animals show strong CO_2_-attraction, opposite to the fed animals. (C) *inx-6(ot808)* mutant L1-arrested animals show reduced CO_2_-attraction. (D) *daf-16(mu86)* mutant L1-arrested animals continue to show CO_2_-attraction, while *daf-16(mu86)* mutant starved adults show CO_2_-avoidance. (E) Expression of fosmid-based *inx-6* reporter transgene, *otIs473[inx-6::sl2::yfp::h2b],* in AIB neurons remains unaffected in *daf-16(mu86)* mutant L1 arrested animals. (F) *daf-2(e979)* mutant Fed-L1s show CO_2_-attraction, as opposed to N2 Fed-L1s. (G) CO_2_-attraction in L1 arrested animals, which were mutant for *daf-2(e979), pdk-1(sa709), akt-1(ok525), akt-2(ok393)* and double-mutant for *age-1(m333); daf-16(m26)* remains unaffected. See also Figure S1.

Our results showed that *C. elegans* larvae at L1-arrest stages also exhibit strong attraction to CO_2_, similar to dauers and starved adult animals **(Figure 1B**, **Table 1).** CO_2_ chemotaxis in adults and dauers depends on BAG neurons and the BAG-expressed receptor-type guanylate cyclase, GCY-9^22,23^. Our data suggested that CO_2_ chemotaxis in L1-arrested animals also depends on both BAG and *gcy-9* **(Figure S1B,C)**. Moreover, *inx-6(ot808)* mutant L1-arrested animals, which eliminates the electrical synapses between BAG and AIB neurons, showed a significant reduction in CO_2_-attraction **(Figure 1C)**, suggesting a conserved role of BAG sensory neurons across developmental stages in *C. elegans*.

**Table 1:** Summary of CO_2_-chemotaxis dependence on distinct regulators at different developmental stages.

| Stage | CO<sub>2</sub> Attraction |  |  | CO<sub>2</sub> Avoidance |  |
| --- | --- | --- | --- | --- | --- |
|  | BAG-AIB gap junction | DAF-16/FOXO | CHN-1/CHIP | IIS Pathway | CRH-1/CREB1 |
| L1 arrest | Partly Dependent | Independent | Dependent |  |  |
| L1 fed |  |  |  | Dependent | Dependent |
| L3 arrest | Not present | Not tested | Dependent |  |  |
| L3 fed |  |  |  | Dependent | Dependent |
| Starved young adult | Not present | Dependent | Dependent |  |  |
| Fed young adult |  |  |  | Not tested | Not tested |
| Dauer | Partly Dependent | Not tested | Independent |  |  |
| Post-dauer |  |  |  | Dependent | Independent |

*C. elegans* larvae can also undergo additional reversible developmental arrests at early L3 and L4 stages, called L3-arrest (L3d) or L4 arrest (L4d), when exposed to food deprivation prior to the L3 and L4 molting checkpoints, respectively ^26^. We found that *C. elegans* larvae at L3-, and L4-arrest stages also exhibit strong attraction to CO_2_ **(Figure 1B**, **Table 1)**. However, in contrast to L1-arrested animals, neither L3-arrested nor starved adult animals display *inx-6* expression in AIB neurons, which is necessary for the BAG-AIB electrical synapse formation **(Figure 1A**, **Table 1)**. These results suggest that equivalent chemosensory responses to CO_2_ across different developmental arrest stages may be regulated by distinct mechanisms.

### CO_2_-attraction in developmental arrested stages and starved adults is independent of the insulin signalling pathway

To understand the underlying molecular mechanisms responsible for CO₂-attraction of *C. elegans* at different developmental arrest stages and in starved adults, we looked into the insulin/IGFR-like signalling (IIS) pathway. Under starved conditions, downregulation of the IIS pathway allows nuclear translocation of the DAF-16/FOXO transcription factor, leading to expression of multiple target genes that are crucial for various physiological and metabolic changes required for L1-, L3-, L4-arrest^9,26^, and dauer diapause. To our surprise, we found that while CO₂-attraction in starved adult animals was dependent on DAF-16 activity, CO₂-attraction in L1-attested animals remained unaffected in *daf-16(mu86)* mutants **(Figure 1D**, **Table 1)**. Similarly, We found that AIB-specific *inx-6* expression in the L1-arrest stage was also unaffected in *daf-16(mu86)* mutant animals, suggesting putative BAG-AIB electrical synapse formation at this stage may be independent of DAF-16/FOXO **(Figure 1E)**. Notably, AIB-specific *inx-6* expression and hence the BAG-AIB electrical synapses formation in dauer animals depend on DAF-16 activity ^19^, further suggesting that stage-specific effectors regulate the CO₂-attraction in different developmental stages.

We next tested whether CO₂-attraction of L1-arrested animals depend on downregulation of the IIS pathway. Notably, the IIS pathway activity was shown to be required for CO₂-attraction in dauers^27^. Downregulation of DAF-2, the sole insulin/IGF receptor in *C. elegans* at the time of hatching of the embryo using a temperature-sensitive allele of *daf-2, daf-2(e979),* was sufficient to induce L1 arrest even in presence of the OP50 diet^9,29^. These ectopic *daf-2(e979)* mutant L1-arrested animals showed strong attraction to CO₂, while non-mutant fed L1 animals continued to show strong CO_2_-avoidance **(Figure 1F)**. This suggested that downregulation of the IIS pathway is sufficient to induce CO₂-attraction in L1-arrested animals. Furthermore, CO₂-attraction in L1-arrested animals remained unaffected in mutants of other downstream members of the insulin pathway, including *age-1(m333)*, *pdk-1(sa709)*, *akt-1(ok525),* and *akt-2(ok393)* **(Figure 1G)**. These results together suggested that CO₂-attraction in L1-arrested animals and dauers is differentially regulated by the IIS pathway.

### Ubiquitin ligase CHN-1/CHIP regulates CO_2_-attraction during developmentally arrested stages and in starved adult *C. elegans*

Next, we wanted to understand the underlying mechanisms responsible for downregulation of the IIS pathway and promotion of CO_2_-attraction in the L1 arrest stage. It was shown that CO_2_-attraction in dauer animals depends on INS-1, an inhibitory insulin-like peptide that antagonizes DAF-2 activity^27^. *ins-1(ot1360)* mutant L1-arrested animals continued to show CO_2_-attraction, although at a significantly reduced level **(Figure S2A)**, suggesting an INS-1-independent mechanism might promote CO_2_-attraction during L1 arrest.

The ubiquitin-ligase CHN-1/CHIP has been shown to negatively regulate DAF-2 activity in L1-arrested and adult animals^12,13^. We hypothesized that CHN-1/CHIP may downregulate DAF-2, and thereby promote CO_2_-attraction in L1-arrested animals. Supporting this hypothesis, *chn-1(by155)* mutant L1-arrested animals exhibited strong CO_2_-avoidance instead of attraction, representing a complete reversal of the CO₂ chemosensory preference **(Figure 2A**, **Table 1)**. Additionally, *chn-1(by155)* mutant L3-arrested animals and starved adults also displayed CO_2_-avoidance, similar to well-fed L3 stage and adult animals, respectively **(Figure 2A**, **Table 1)**. This effect of *chn-1/CHIP* mutation was specific to CO_2_-chemotaxis, as *chn-1(by155)* mutant animals maintained attraction to diacetyl, and avoidance to 2-octanone **(Figure S2B,C)**. To determine whether CHN-1/CHIP regulates CO_2_-chemotaxis by modulating DAF-2 functions, we tested the CO_2_-chemotaxis of *chn-1(by155)*; *daf-2(e979)* double mutant animals. Consistent with the hypothesis, *chn-1(by155)*; *daf-2(e979)* double mutant L1-arrested animals showed CO_2_-attraction, in contrast to *chn-1(by155)* mutants alone, suggesting *daf-2* is epistatic to *chn-1/CHIP* **(Figure 2B)**.

**Figure 2:**
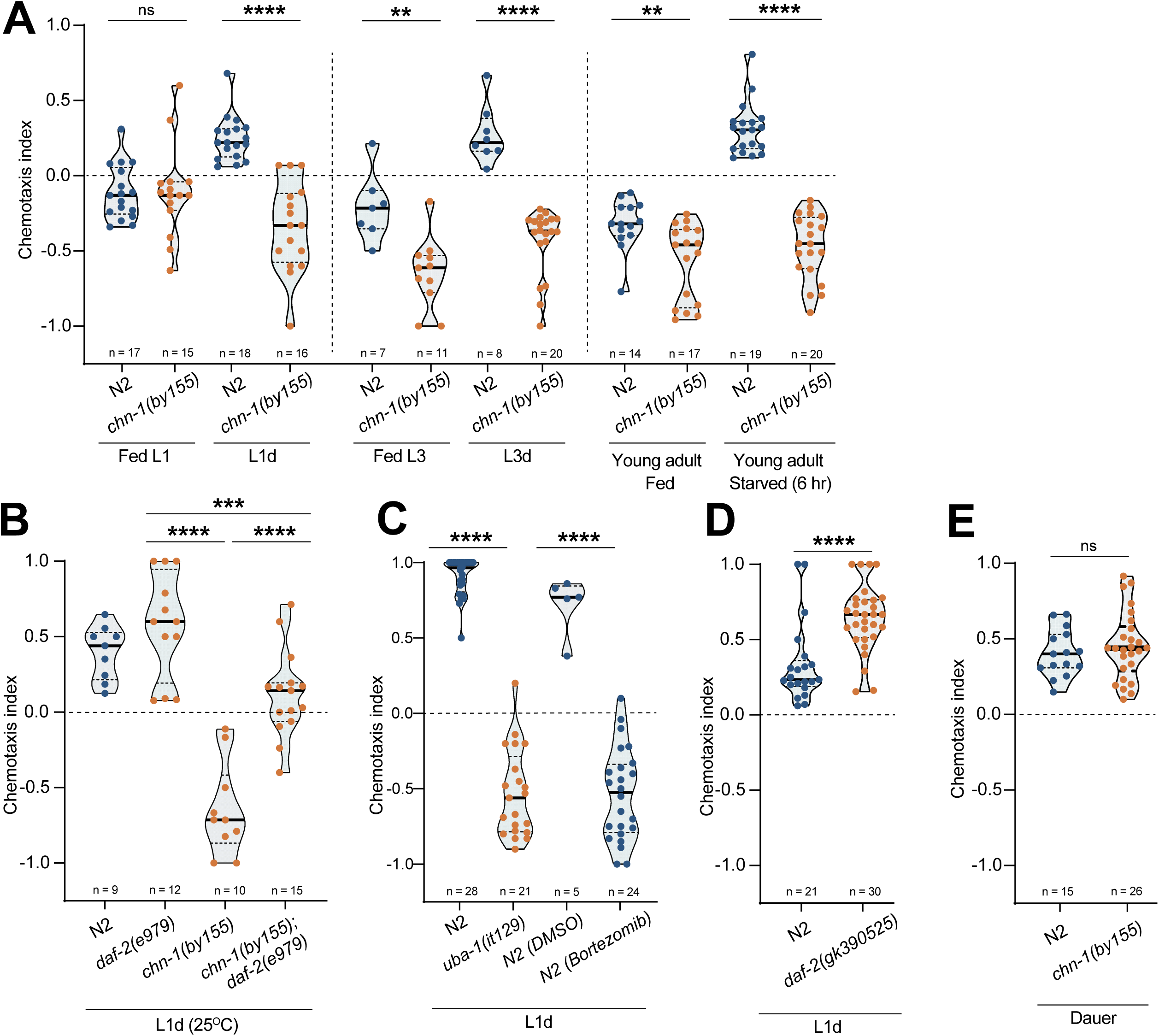
CHN-1/CHIP is required for CO_2_-attraction in L1-, L3-arrested, and starved young adult animals. (A-E) Each circle represents the chemotaxis index calculated from a single chemotaxis assay. Violin plots represent the distribution, black horizontal solid lines represent the median and horizontal dotted lines represent the quartiles. Mann–Whitney U tests p-values for each comparison: n.s., nonsignificant, *p < 0.05, **p < 0.01, ***p < 0.001, ****p < 0.0001. (A) *chn-1(by155)* mutant L1- and L3-arrested, and starved young adult animals show strong CO_2_-avoidance, opposite to matched N2 control animals, which show CO_2_-attraction. *chn-1(by155)* mutant Fed-L1, L3 and young adult animals continue to show CO_2_-avoidance. (B) CO_2_-avoidance of *chn-1(by155)* mutant L1-arrested animals is reversed by *daf-2(e979)* mutation. (C) L1-arrested animals, which are mutant for *uba-1(it129)* or treated with Bortezomib, show strong CO_2_-avoidance, opposite to DMSO treated control L1-arrested N2 animals, which show CO_2_-attraction. (D) *daf-2(gk390525)* mutation does not affect CO_2_-attraction in L1-arrested animals. (E) *chn-1(by155)* mutation does not affect CO_2_-attraction in dauers. See also Figure S2.

To determine whether the CHN-1/CHIP regulates CO_2_-chemotaxis in L1-arrested animals via ubiquitination, we tested CO_2_-chemotaxis in animals mutant for *uba-1*, the sole E1-activating enzyme encoded in the *C. elegans* genome^30^. Similar to *chn-1/CHIP* mutant and fed-L1 animals, *uba-1(it129)* mutant L1-arrested animals also showed strong CO_2_-avoidance **(Figure 2C)**, further supporting the role of CHN-1/CHIP-mediated ubiquitination in regulating CO_2_-attraction during the L1-arrest stage. In adult *C. elegans,* CHN-1/CHIP inhibits DAF-2 activity independent of proteasomal degradation through mono-ubiquitinating DAF-2 at lysine, K1614, thereby regulating proteostasis and aging^12^. This regulation is disrupted in *daf-2(gk390525)* mutant animals harbouring the K1614E mutation^12^. We found that CO_2_-attraction in *daf-2(gk390525)* L1-arrested animals remained unaffected **(Figure 2D)**, suggesting that K1614-independent regulation of DAF-2 by CHN-1/CHIP may control CO_2_-chemotaxis valence during L1-arrest stage. Furthermore, inhibition of proteasomal degradation in L1-arrested animals using Bortezomib, an inhibitor of the large subunit of proteasome, was sufficient to reverse CO_2_-attraction to avoidance, similar to *uba-1* and *chn-1/CHIP* mutant animals **(Figure 2C).** These results together suggest an L1-arrest stage-specific regulation of DAF-2 by CHN-1/CHIP that modulates CO_2_-chemotaxis during this developmental stage.

We next tested whether CO_2_-attraction in dauer animals also depend on CHN-1/CHIP or ubiquitin-proteasome pathway. In contrast to developmentally arrested and starved adult animals, CO_2_-attraction remained unaffected in *chn-1(by155)* and *uba-1(it129)* mutant dauers **(Figure 2E, S2D, Table 1)**. These results together suggested that CO₂-attraction in animals at different developmental stages, including L1-arrest and dauer diapause, depends on distinct regulatory mechanisms.

### Insulin/IGF-1 signalling promotes rapid reversal of CO_2_-attraction upon feeding

Previously, it was shown that CO_2_-attraction in long-term starved adults can be reversed upon two hours of feeding on *E.Coli* OP50^23^. Surprisingly, we found that L1-arrested animals displaying strong CO_2_-attraction undergo a rapid reversal of CO_2_-chemotaxis valence from attraction to avoidance within 15 minutes of feeding on *E.Coli* OP50 **(Figure 3A)**. Although exposure to bacterial conditioned LB media alone significantly reduced CO_2_-attraction in L1-arrested animals, it was not sufficient to induce a complete reversal to CO_2_-avoidance **(Figure S3A)**, suggesting that feeding on bacterial food is required for the reversal of CO_2_-chemotaxis valence during L1-arrest exit. On the other hand, dauers undergo reversal CO_2_-chemotaxis valence to avoidance only after 16 hours of feeding on *E.Coli* OP50 **(Figure 3B)**. These observations prompted us to investigate the molecular mechanisms regulating the reversal of CO_2_-attraction in two distinct developmental stages, L1-arrest and dauer diapause.

**Figure 3:**
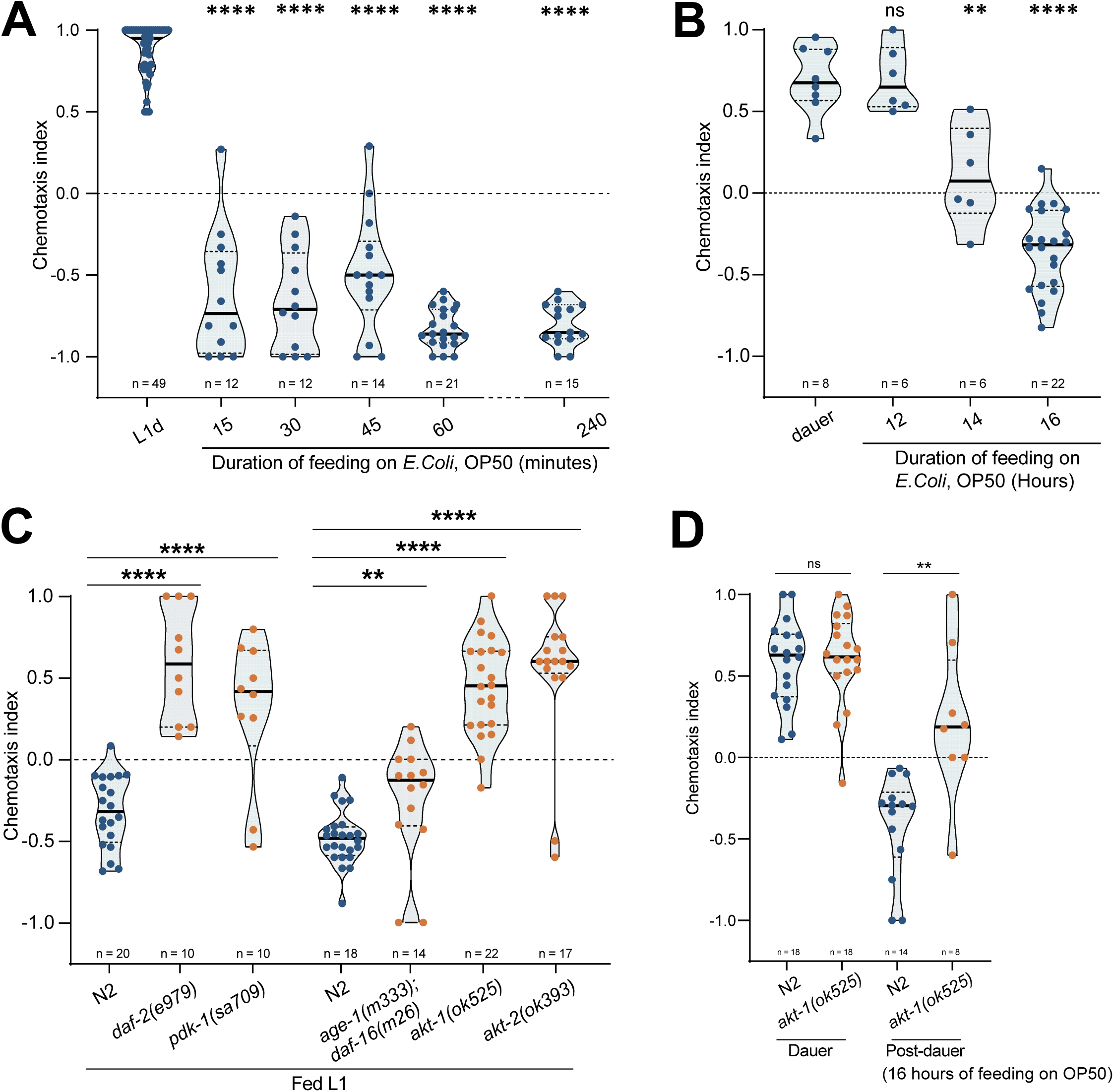
IIS pathway and CRH-1/CREB1 differentially regulate feeding-dependent reversal of CO_2_-attraction in distinct life stages. Each circle represents the chemotaxis index calculated from a single chemotaxis assay. Violin plots represent the distribution, black horizontal solid lines represent the median and horizontal dotted lines represent the quartiles. Mann–Whitney U tests p values for each comparison: n.s., nonsignificant, *p < 0.05, **p < 0.01, ***p < 0.001, ****p < 0.0001. (A) Feeding on *E. coli* OP50 can reverse CO_2_-attraction of L1-arrested animals within 15 minutes. (B) Reversal of CO_2_-attraction of dauer animals requires feeding on *E. coli* OP50 for 16 hours (C) *daf-2(e979), pdk-1(sa709), akt-1(ok525)* and *akt-2(ok393)* mutant fed-L1 animals show CO_2_-attraction, opposite to N2 Fed-L1 animals, which show CO_2_-avoidance. *age-1(m333); daf-16(m26)* double-mutant fed L1 animals show weak CO_2_-avoidance. (D) *akt-1(ok525)* mutant dauers fail to induce CO_2_-avoidance upon feeding. See also Figure S3.

Since CHN-1-mediated downregulation of the IIS pathway is required for CO_2_-attraction during the L1-arrest stage, we tested the role of the IIS pathway in reversal of the chemotaxis valence during L1-arrest exit. Our data showed that during the L1-arrest exit, feeding-induced reversal of CO_2_-chemotaxis to avoidance depends on the activity of the IIS pathway, including DAF-2, and downstream kinases, AGE-1, PDK-1, AKT-1, and AKT-2 **(Figure 3C**, **Table 1)**. Similarly, reversal of CO_2_-chemotaxis during dauer exit also required activity of AKT-1 **(Figure 3D**, **Table 1)**, suggesting conserved roles of the IIS pathway in reversal of CO_2_-chemotaxis upon feeding across developmental stages.

### Activity of CRH-1a/CREB1 isoform in the BAG sensory neuron is required to promote CO_2_-avoidance upon feeding, during L1-arrest exit

As CO_2_-attraction in L1-arrested animals was independent of DAF-16/FOXO activity, we hypothesized that the IIS pathway may promote CO2-avoidance during L1-arrest exit through regulating a different downstream transcription factor. In a screen to identify potential regulators of feeding-dependent reversal of CO_2_-chemotaxis valence during L1-arrest exit, we identified the conserved transcription factor CRH-1, a homolog of mammalian cAMP response element-binding protein (CREB). CRH-1/CREB1 has been shown to be regulated by a multitude of signalling pathways in both vertebrates and invertebrates, including the IIS pathway^31,32^. The vertebrate homolog of the *C. elegans* AKT-1, Akt1/PKB, has been shown to directly activate CREB1^33^. We found that L1-arrested animals mutant for *crh-1(tz2)* failed to reverse CO_2_-chemotaxis valence upon feeding on E.coli OP50 for 1 hour, and continued to show strong CO_2_-attraction **(Figure 4A**, **Table 1)**. Moreover, *crh-1(tz2)* mutant L3-arrested animals also failed to reverse CO_2_-chemotaxis valence upon feeding **(Figure S4A, Table 1),** as well as, well-fed, L3-stage animals that never transited through developmental arrest stages also showed strong CO_2_-attraction, unlike control well-fed L3-stage animals **(Figure S4B)**.

**Figure 4:**
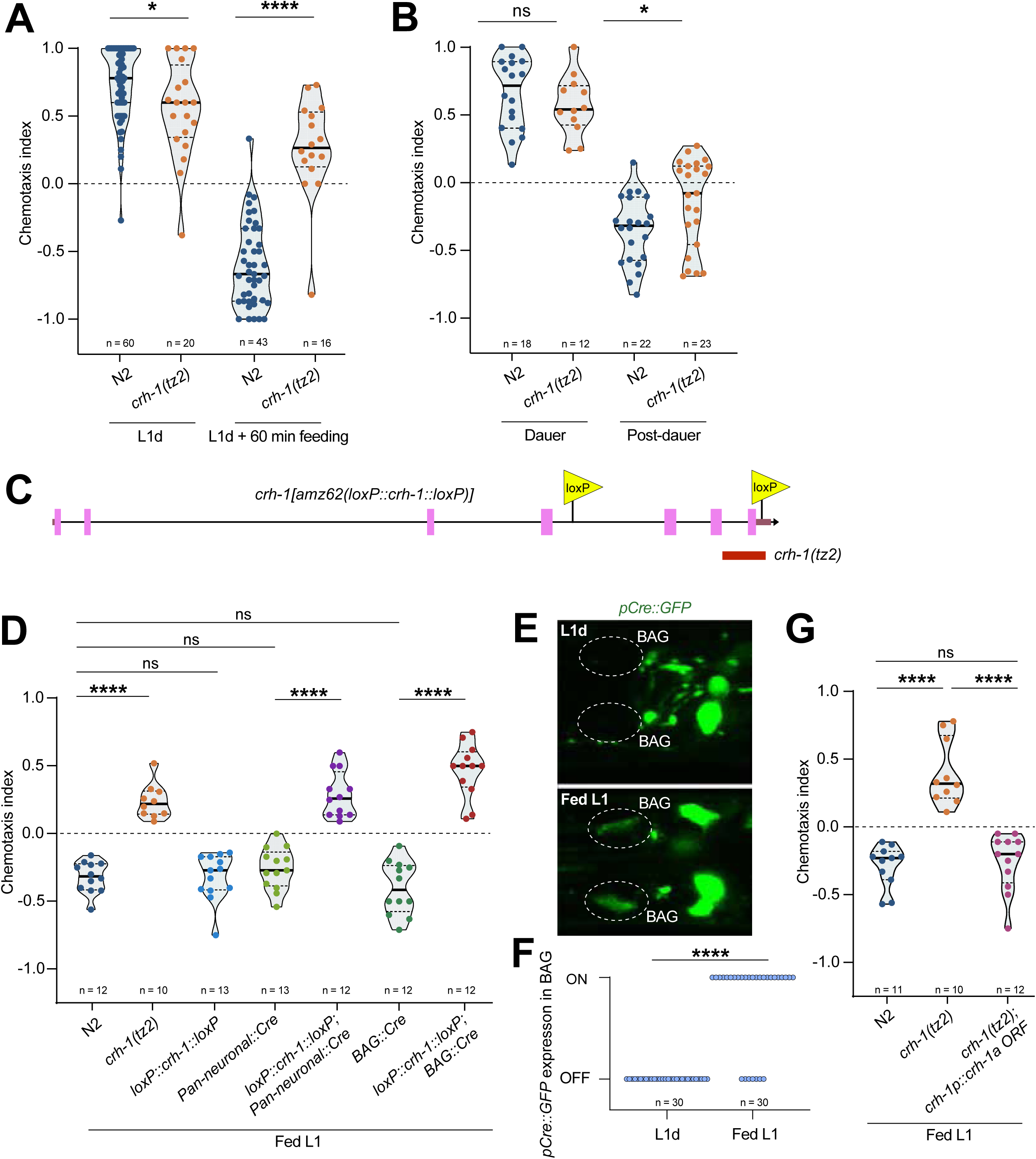
CRH-1/CREB1 functions in BAG neurons to promote CO_2_-avoidance during L1-arrest exit. (A, B, D and G) Each circle represents the chemotaxis index calculated from a single chemotaxis assay. Violin plots represent the distribution, black horizontal solid lines represent the median and horizontal dotted lines represent the quartiles. Mann–Whitney U tests p values for each comparison: n.s., nonsignificant, *p < 0.05, **p < 0.01, ***p < 0.001, ****p < 0.0001. (A) *crh-1(tz2)* mutant fed-L1 animals show CO_2_-attraction, opposite to N2 Fed-L1 animals, which show CO_2_-avoidance. (B) *crh-1(tz2)* mutation does not affect CO_2_-avoidance in post-dauer animals. (C) Schematic of the *C. elegans crh-1* locus showing positions of loxP recombination sites in the *crh-1(amz62[loxP::crh-1::loxP])* allele. (D) Cre-recombinase mediated pan-neuronal or BAG-specific deletion of *crh-1* in *loxP* flanked *crh-1* animals prevented feeding dependent reversal of CO_2_ chemotaxis to avoidance, similar to fed *crh-1(tz2)* mutant L1 animals. (E) *pCre::GFP* reporter gets upregulated in BAG neurons during L1-arrest exit, upon feeding for 2 hours on *E. Coli* OP50. (F) Quantification of data shown in panel 4E. Mann–Whitney U tests, ****p < 0.0001. (G) Loss of CO_2_-avoidance phenotype in *crh-1(tz2)* mutant fed L1 animals can be rescued by expressing *crh-1a* isoform.

Interestingly, in contrast to the post-L1-arrest animals, *crh-1(tz2)* mutant post-dauer animals that were fed for 16 hours on *E.Coli* OP50 showed CO_2_-avoidance **(Figure 4B**, **Table 1)**, suggesting CRH-1/CREB1 differentially regulates feeding-induced reversal of CO_2_-attraction during L1-arrest and dauer diapause stage exit. CRH-1/CREB1 has been shown to function in various neuron-types to regulate a multitude of physiological processes, including aging, lifespan, chemotaxis behaviour, and associative memory formation ^16,17,34,35^. To test where CRH-1/CREB1 functions to promote CO_2_-avoidance upon feeding, a loxP recombination-site-flanked conditional knockout allele of *crh-1*, *crh-1(amz62[loxP::crh-1::loxP])*, was engineered **(Figure 4C)**. In the presence of the Cre recombinase, this conditional allele deleted a 3.5 kb genomic region encompassing the entire *crh-1f* coding sequence and the conserved last three exons of all other *crh-1* isoforms, *crh-1(a-e* and *g)* **(Figure 4C)**. Our data suggested that loss of *crh-1* specifically in the nervous system using pan-neuronal expression of the Cre recombinase was sufficient to inhibit feeding-induced onset of CO_2_-avoidance during L1-arrest exit **(Figure 4D)**. Finally, we show that loss of *crh-1/CREB1* specifically in the BAG sensory neurons was sufficient to abrogate feeding-dependent reversal of CO_2_-chemotaxis valence **(Figure 4D)**. Supporting these findings, we found that the expression of a CRH-1/CREB1-reporter transgene, *pCRE::gfp^36^,* where GFP expression is controlled by CRH-1/CREB1 response element, was upregulated in BAG neurons upon feeding during L1-arrest exit **(Figure 4E,F)**.

The *crh-1* locus encodes seven different splice isoforms (*crh-1 a-g*) that regulate diverse functions, including stress responses and memory formation^17,37-39^. Of these, six isoforms, *crh-1(a-e* and *g),* encode both the kinase-activated domain and the DNA-binding bZIP domain. Expression of the *crh-1a* isoform, using a 3kb cis-regulatory region upstream of the *crh-1* locus, completely rescued the CO_2_-chemotaxis defect observed in *crh-1(tz2)* mutant L1-stage animals **(Figure 4G)**. These results suggested that CRH-1a isoform functions at the sensory neuron level to promote feeding-induced reversal of CO_2_-chemotaxis valence during L1-arrest exit.

## DISCUSSION

Organisms can sense internal physiological states and modify behavioural response to external cues, supporting growth, metabolism, food and mate searching, and survival. In this study we show that, in addition to previously described dauer animals and long-term starved adults, *C. elegans* larvae at other developmental checkpoint arrest stages (L1-, L3-, and L4-arrests) also exhibit CO₂-attraction by alteration in chemotaxis valence. Plasticity in CO₂-chemotaxis in dauer-equivalent stages is conserved among nematode phyla and plays a crucial role in the life cycle of parasitic nematodes by acting as a host seeking mechanism^22,40^. Further work will be needed to determine conservation of the CO₂-chemotaxis plasticity in other developmental arrest stages. Previously studies have shown that distinct neural circuits involving different interneurons regulate CO₂-attraction in dauers and long-term starved adults^28^. We show here that development clock of *C. elegans* determines the molecular mechanisms by which chemosensory responses to environmental CO_2_ concentrations are altered under different nutritional conditions. We show, even at the level of CO_2_-sensing BAG neurons, distinct molecular mechanisms at different life stages regulate the chemosensory response to CO_2_.

Our findings indicate that the conserved quality-control ubiquitin ligase CHN-1/CHIP regulates the insulin/IGF receptor DAF-2 through ubiquitination and proteasomal degradation pathway, promoting CO₂-attraction in the L1-arrest stage and in starved adults, but not during the dauer diapause **(Figure 5)**. This result is also consistent with published data showing that CO_2_-attraction in dauers depends on DAF-2 and IIS pathway activity^27^. Additionally, CHN-1/CHIP mono-ubiquitinates DAF-2 at lysine K1614 in adults, but not during the L1-arrest stage^12,41^. We show here that CHN-1/CHIP regulates CO₂-attraction in the L1-arrest stage independently of K1614. A previous work has shown that a dimer-monomer switch controls substrate selection and ubiquitination by CHN-1/CHIP^42^. This work provides a foundation for understanding how stage-specific developmental programs regulate the interaction between CHN-1/CHIP and DAF-2, which may have broader implications for development, metabolic regulation, and aging. Furthermore, CHN-1/CHIP-mediated downregulation of DAF-2 regulates CO₂-attraction through distinct downstream effectors at different life stages. While DAF-16/FOXO activity is required in starved adults, CO₂-attraction in the L1-arrest stage is independent of DAF-16/FOXO **(Figure 5)**, suggesting the involvement of a distinct L1-stage-specific effector in regulating CO_2_-attraction and BAG-AIB gap junction formation.

**Figure 5:**
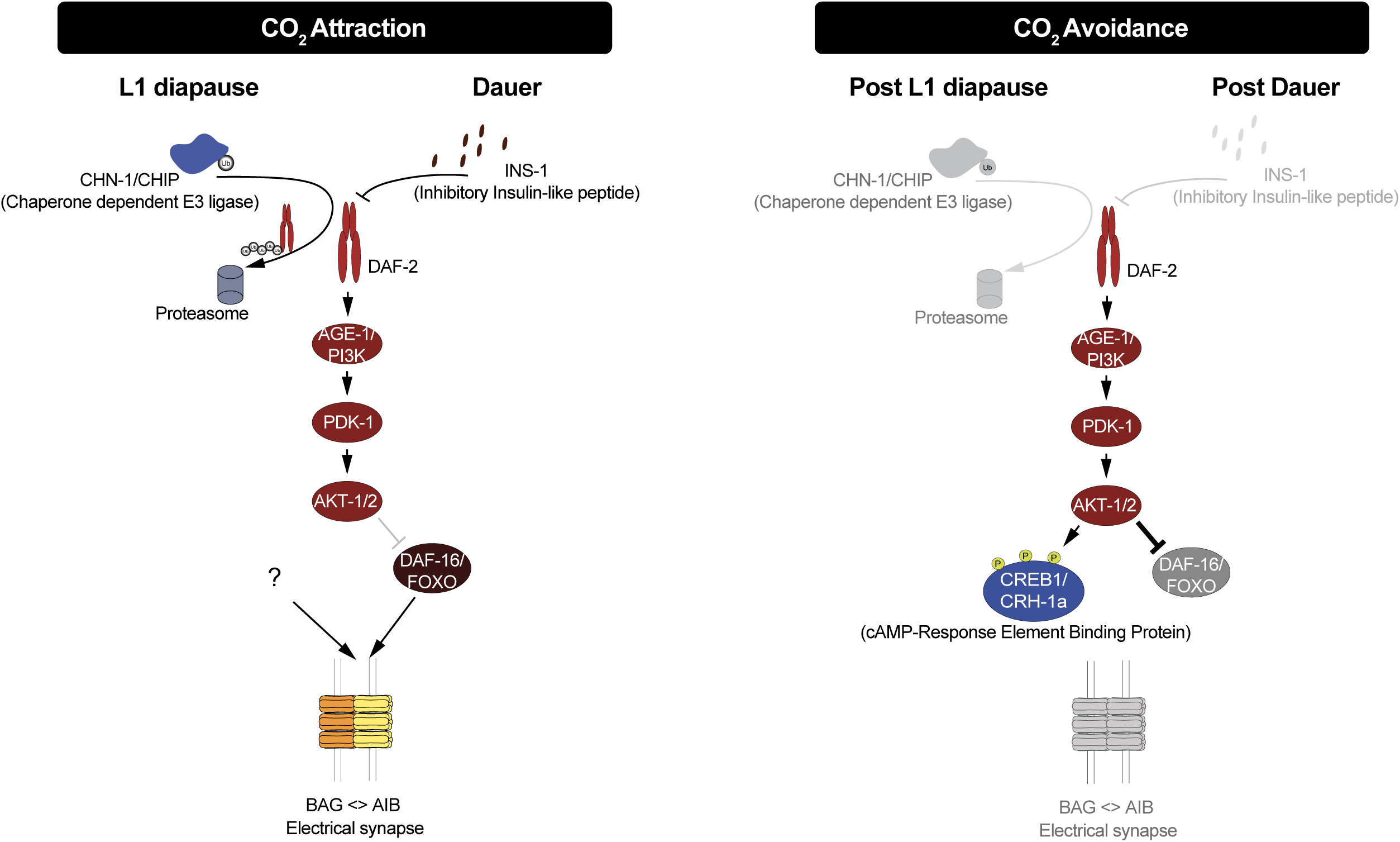
Distinct molecular mechanisms regulate plasticity of CO_2_-chemotaxis valence at different life stages. Schematic showing distinct molecular mechanisms regulate insulin/IGF signalling receptor DAF-2 in L1 arrested and dauer animals to promote CO_2_-attraction. INX-6 expression in AIB neurons and the resultant BAG-AIB electrical synapse formation in L1 arrested and in dauer animals is also mediated by distinct molecular mechanisms. Similarly, CO_2_-avoidance upon feeding is also promoted by distinct, stage-specific molecular mechanisms.

Our work indicates that feeding-dependent upregulation of the IIS pathway sculpts the CO_2_-chemotaxis behaviour by promoting avoidance. Additionally, we identify here that the CRH-1/CREB1 transcription factor functions cell autonomously in the BAG sensory neuron to regulate reversal of the CO_2_-chemotaxis upon feeding during developmental checkpoint stages, but not during the alternative dauer diapause stage **(Figure 5)**. Moreover, we show that CRH-1/CREB1 is also continuously required to maintain CO_2_-avoidance during the L3 stage, which nematodes bypass to enter the dauer stage. These results provide a framework for understanding how developmental programs impinge on the CRH-1/CREB1 pathway to modulate chemosensory behaviour at specific life stages. Additionally, this work also provides a foundation for further study into how CRH-1/CREB1 regulates CO_2_-chemosensory behaviour in parasitic nematodes, which utilize BAG-equivalent neurons to detect CO_2_ as a host cue and modulate their CO_2_-chemotaxis behaviour during different life stages to infect hosts^22,40^.

## EXPERIMENTAL MODEL AND SUBJECT DETAILS

### *C. elegans* strains and handling

*C. elegans* strains cultured on NGM plates that are coated with *E. coli* (OP50) bacteria as a food source, as per the standardized protocols unless otherwise mentioned^43,44^. Strains were maintained at 20-22°C. All temperature sensitive alleles were maintained at 15°C. *C. elegans* strain N2-Bristol was used as wild-type control. Apart from *daf-2* mutant animals, dauer arrest was induced under standard starvation, crowding and high-temperature conditions, as described previously^19^. Dauer animals were selected by 1% SDS treatment from the lid of the plates. Transgenic strains were generated using microinjection technique. Detailed injection conditions are indicated in the strain list (Table S2).

## METHOD DETAIL

### CO_2_ Chemotaxis assay

CO_2_ chemotaxis assays for animals of all stages were performed on standard 9cm NGM plates. All CO_2_-chemotaxis assays were performed as previously described ^19,22^. CO_2_ gradient on the NGM plates was generated by pumping a mixture of (10% CO2 + 20% O2 + 70% N2) from an inlet and a mixture of (20% O2 + 80% N2) from another inlet (as indicated in Figure S1), using programable syringe pumps (New Era Scientific Pump Systems) at flow rates of 1.5ml/min for non-dauer assays and 0.5ml/min for dauer assays.∼100-150 animals were placed at the centre of the assay plates. After 30 minutes, the number of animals inside the air and CO_2_ scoring regions [2cm circles under respective gas inlets], were counted. The chemotaxis index (C.I.) was calculated as:

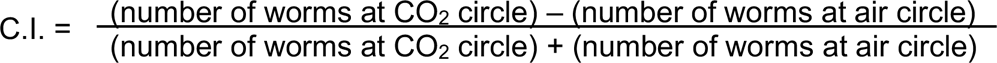

CO_2_-chemotaxis assays involving insulin/IGF-1 pathway mutant L1 stage animals (excluding *daf-16* and *daf-16; age-1* and *daf-2(gk390525)* mutants) were performed with 0.5ml/min flow rate for 90 minutes, along with control genotypes. CO_2_-chemotaxis assays involving insulin/IGF-1 pathway mutant dauer stage animals (excluding *daf-16* mutants) were performed with 0.5ml/min flow rate for 30 minutes, along with control genotypes. Assays involving *crh-1* knockout and rescue lines were performed with 0.5ml/min flow rate for 60 minutes. CO_2_-chemotaxis assays of L1 larval stage and young adults for *chn-1, daf-2(e979)* and *chn-1;daf-2* double mutants were performed at 1.5ml/min flow rate for 30 minutes. For these assays, animals that had moved beyond 1 cm line towards either the air or CO₂ side (i.e. beyond the blue lines indicated in the Figure S1) were counted and the chemotaxis index was subsequently calculated by using the above formula.

Synchronised L1-arrested population was prepared by hatching isolated embryos in M9 buffer overnight^44^. This population was habituated on NGM agar plates for 2 hour prior to chemotaxis assay. For fed-L1 population, habituated L1-arrested animals were allowed to feed on E.Coli OP50 on NGM agar plates at 25°C for desired length of time. L3- and L4-arrested animals were isolated as described previously ^26^. In brief, synchronised well-fed population was prepared by hatching isolated embryos in M9 buffer overnight to get L1-arrested population. This population was fed with E.Coli OP50 on NGM agar plates at 25°C with to get synchronised well-fed population. L3- and L4-arrested animals were isolated by starving a well-fed synchronised population at mid-L2 and mid-L3 larval stages, respectively, for 24 hours at 25°C. The stage was further confirmed by checking the vulva morphology prior to chemotaxis assays. Starved adult animals were prepared by selecting well fed 1 day old adult hermaphrodites and starving them for 6 hours without food on NGM plates. Well fed, continuous developing L3 stage animals were obtained by hatching isolated embryos on NGM agar plates supplemented with E.Coli OP50. For CO_2_-chemotaxis assays involving temperature sensitive alleles of *uba-1, daf-2, pdk-1,* synchronised L1-arrested population were cultivated for additional 24 hours at 25°C on NGM agar plates prior to chemotaxis assays.

### Bortezomib treatment

Synchronised L1-arrested population was incubated 500ul of S-Basal medium^44^ supplemented with 40uM Bortezomib (Sigma-Aldrich, Cat. Number 504314) in a 2ml microcentrifuge tube on a nutator for overnight, followed by habituated on NGM agar plates supplemented with 40uM Bortezomib for 2 hours prior to chemotaxis assay. Control groups were treated similarly with DMSO solvent. CO_2_-chemotaxis assays were performed on a 90cm NGM agar plate with 0.5ml/min gas flow rate for 1hour.

### CRISPR/Cas9-mediated genome editing

Floxed-alleles of *crh-1* were generated using purified Cas9 (IDT, catalog # 1081059), tracrRNA (IDT, catalog # 1072533) and crRNA (IDT) based on previously published protocols ^45,46^. We inserted two *loxP* sites flanking the *crh-1* genomic locus: 5’ loxP within the 4^th^ intron; crRNA sequence 5’ ctactcagcacagttaaagg 3’ 3’ loxP within the 3’ UTR; crRNA sequence 5’ gtagatatccattaagtggg 3’ **(Figure 4C)**.

### Generation of Cre recombinase constructs

Sequences of a synthetic pan-neuronal driver (*UPNp)* driver^47^, and BAG-specific *flp-17p* driver were amplified using PCR and cloned into a plasmid containing *3XNLS::Cre*::*unc-54 3’ UTR* cassette in the pPD95.75 vector backbone, using SphI and SmaI restriction sites.

### Generation of *CREB/crh-1* rescue construct

For amplification of the *crh-1a* ORF, total cDNA was synthesized using oligo(dT)_20_ primer and SuperScript III First-Strand Synthesis System (Thermo Fisher Scientific; Catalog #18080051), according to the standard protocol. ORF of *crh-1a* was amplified using gene-specific primer, and cloned into pMiniT2.0 vector (NEB). This ORF was fused to a *sl2::2xNLS::tagrfp::unc-54 3’ UTR* cassette in the miniMOS plasmid backbone^48^. To generate *crh-1p::crh-1a::sl2::2xNLS::tagrfp*, a 3.0kb cis-regulatory region (-3017 to 0) was ligated upstream of the *crh-1a::sl2::2xNLS::tagrfp* cassette, using SmaI and restriction sites.

### Microscopy

Worms were anesthetized using 100mM of sodium azide and mounted on 5% agarose pads on glass slides. Imaging were done using either Zeiss 880 or NIKON AX confocal laser-scanning microscopes. Image processing and analysis was done by scanning the full Z stack using NIH Fiji software. Maximum intensity projections of representative images were constructed using NIH Fiji software. Figures were prepared using Adobe Photoshop 2025 and Adobe Illustrator 2025. Separate channels were usually adjusted independently using Levels and Curves in Adobe Photoshop.

### Quantification and Statistical Analysis

Statistical tests were selected based on appropriate data type, distribution, and sample size. Across all statistical tests, p<0.05 was considered significant.

For all chemotaxis assays, Shapiro–Wilk tests were first performed to assess normality of the data distribution. For normally distributed data points, the Unpaired t-test (two-tailed) was used for comparing means. For data sets that were not normally distributed and involved two-group comparisons, the Mann–Whitney U test (two-tailed) was used. All the statistical tests and plots were made using GraphPad Prism.

## CONTACT FOR REAGENT AND RESOURCE SHARING

Request for further information, resources and reagents will be fulfilled by the corresponding author, Abhishek Bhattacharya.

## ACKNOWLEDGMENTS

We thank Oliver Hobert as initial experiments were performed in his laboratory; Niels Ringstad and Jung-Hwan Choi for helping with the CO_2_ chemotaxis assay, Pallab Majee, Nargis Mustaq and Bedant Barik for their help with the standardization of experiments in AB lab; Selvanayaki Eswaramoorthy and NCBS *C. elegans* facility for assistance with worm experiments; wormbase.org and wormwiring.org for resources; Caenorhabditis Genetics Center (CGC), Arnab Mukhopadhyay, Kavita Babu, Niels Ringstad, Yoshishige Kimura for *C. elegans* strains; Kavita Babu, Oliver Hobert, Sandhya Koushika, Raghu Padinjat, Urmi Bandyopadyay and members of the Bhattacharya lab for comments on this manuscript. This work was supported by DBT Wellcome Trust India Alliance (IA/I/20/2/505211) and intramural funds from National Centre for Biological Sciences – Tata Institute of Fundamental Research.

## AUTHOR CONTRIBUTIONS

A.R.S., S.M. and A.B. designed experiments and oversaw the project, A.R.S. generated most of the strains and performed most of the experiments, S.M. performed chemotaxis experiments with the insulin/IGF-1pathways genes, A.V. generated the pan-neuronal Cre recombinase expressing construct and the transgenic line. A.R.S., S.M. and A.B. analysed and interpreted data. A.R.S. and A.B. wrote the paper.

## SUPPLEMENTARY FIGURE LEGENDS

**Figure S1: BAG sensory neurons and GCY-9 are required for CO_2_-aattaction in L1-arrested animals**

(A) Schematic of CO_2_-chemotaxis assay. Different scoring regions are marked (described in details in Methods)

(B-C) Each circle represents the chemotaxis index calculated from a single chemotaxis assay. Violin plots represent the distribution, black horizontal solid lines represent the median and horizontal dotted lines represent the quartiles. Mann–Whitney U tests p values for each comparison: n.s., nonsignificant, *p < 0.05, **p < 0.01, ***p < 0.001, ****p < 0.0001.

(B) *ets-5(tm1794)* mutant L1-arrested or fed L1 animals that lack BAG neurons do not chemotax in CO_2_ gradient.

(C) *gcy-9(tm2816)* mutant L1-arrested or fed L1 animals do not chemotax in CO_2_ gradient.

**Figure S2: *chn-1* mutation does not affect chemotaxis in general**

(A-D) Each circle represents the chemotaxis index calculated from a single chemotaxis assay. Violin plots represent the distribution, black horizontal solid lines represent the median and horizontal dotted lines represent the quartiles. Mann–Whitney U tests p values for each comparison: n.s., nonsignificant, *p < 0.05, **p < 0.01, ***p < 0.001, ****p < 0.0001.

(A) *ins-1(ot1360)* mutant L1-arrested animals continued to show CO_2_-attraction, although at a reduced level.

(B-C) Mutation in *crh-1(tz2)* does not affect chemotactic response to 2-Octanone and Diacetyl.

(D) *uba-1(it129)* mutant dauers continued to show CO_2_-attraction.

**Figure S3: IIS pathway and CRH-1 affects feding-dependent reversal of CO_2_-chemotaxis in specifc life stages**

Each circle represents the chemotaxis index calculated from a single chemotaxis assay. Violin plots represent the distribution, black horizontal solid lines represent the median and horizontal dotted lines represent the quartiles. Mann–Whitney U tests p values for each comparison: n.s., nonsignificant, *p < 0.05, **p < 0.01, ***p < 0.001, ****p < 0.0001.

(A) Complete reversal of CO_2_-chemotaxis valence to avoidance requires feeding on *E. coli* OP50.

**Figure S4: CRH-1 affects feeding-dependent reversal of CO_2_-chemotaxis in L3 stage**

Each circle represents the chemotaxis index calculated from a single chemotaxis assay. Violin plots represent the distribution, black horizontal solid lines represent the median and horizontal dotted lines represent the quartiles. Mann–Whitney U tests p values for each comparison: n.s., nonsignificant, *p < 0.05, **p < 0.01, ***p < 0.001, ****p < 0.0001.

(B) *crh-1(tz2)* mutant L3-arrested animals fail to induce CO_2_-avoidance upon feeding.

(C) *crh-1(tz2)* continuously fed L3-stage animals exhibit CO_2_-attarction, while fed L3-stage N2 animals exhibit CO_2_-avoidance.

