## Supplementary figures and images for "Distinct Molecular Mechanisms Regulate Feeding State-Dependent CO_2_ Chemotaxis Plasticity During Different Life Stages in *Caenorhabditis elegans*"

### Supplemental Figures S1 - S4

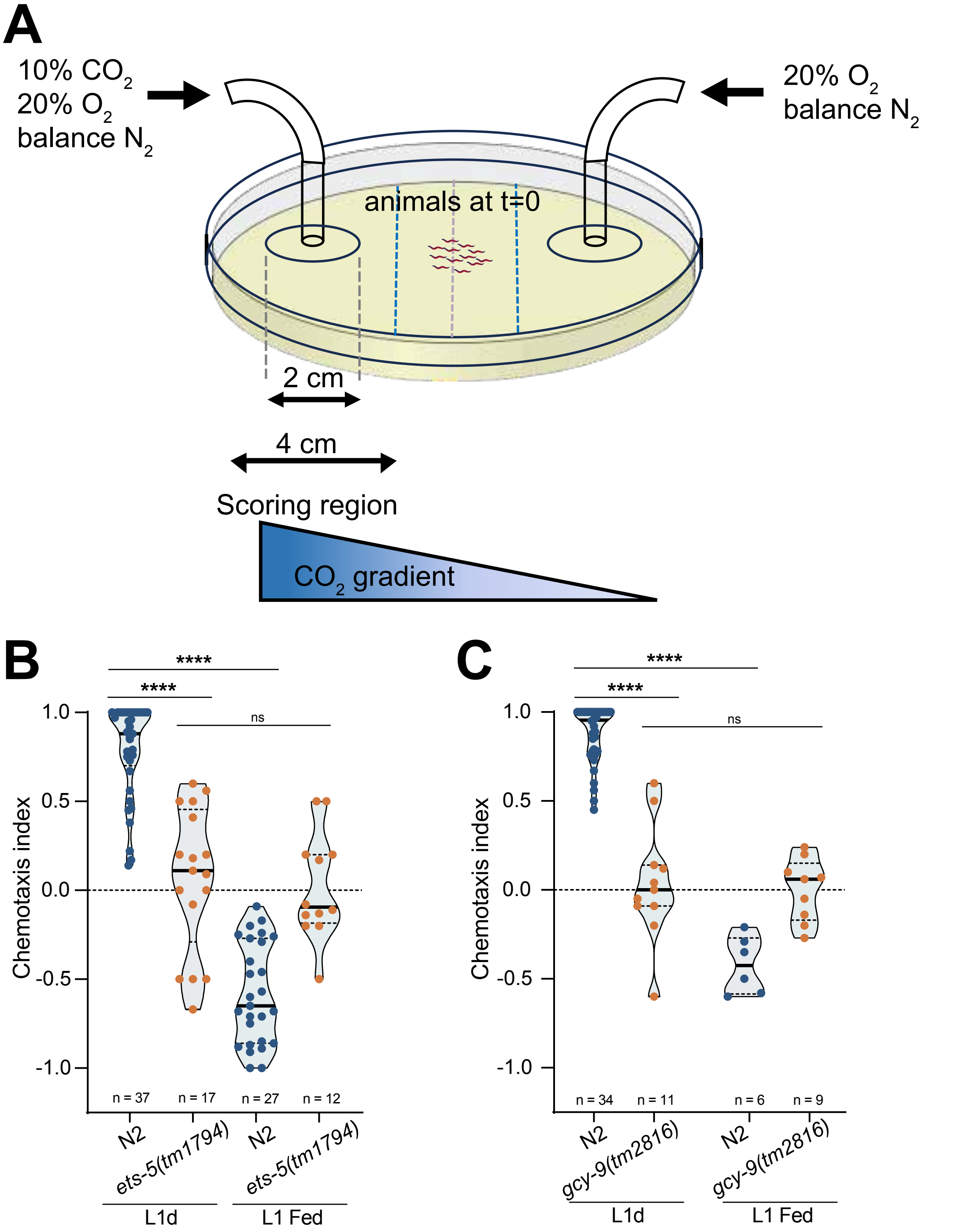

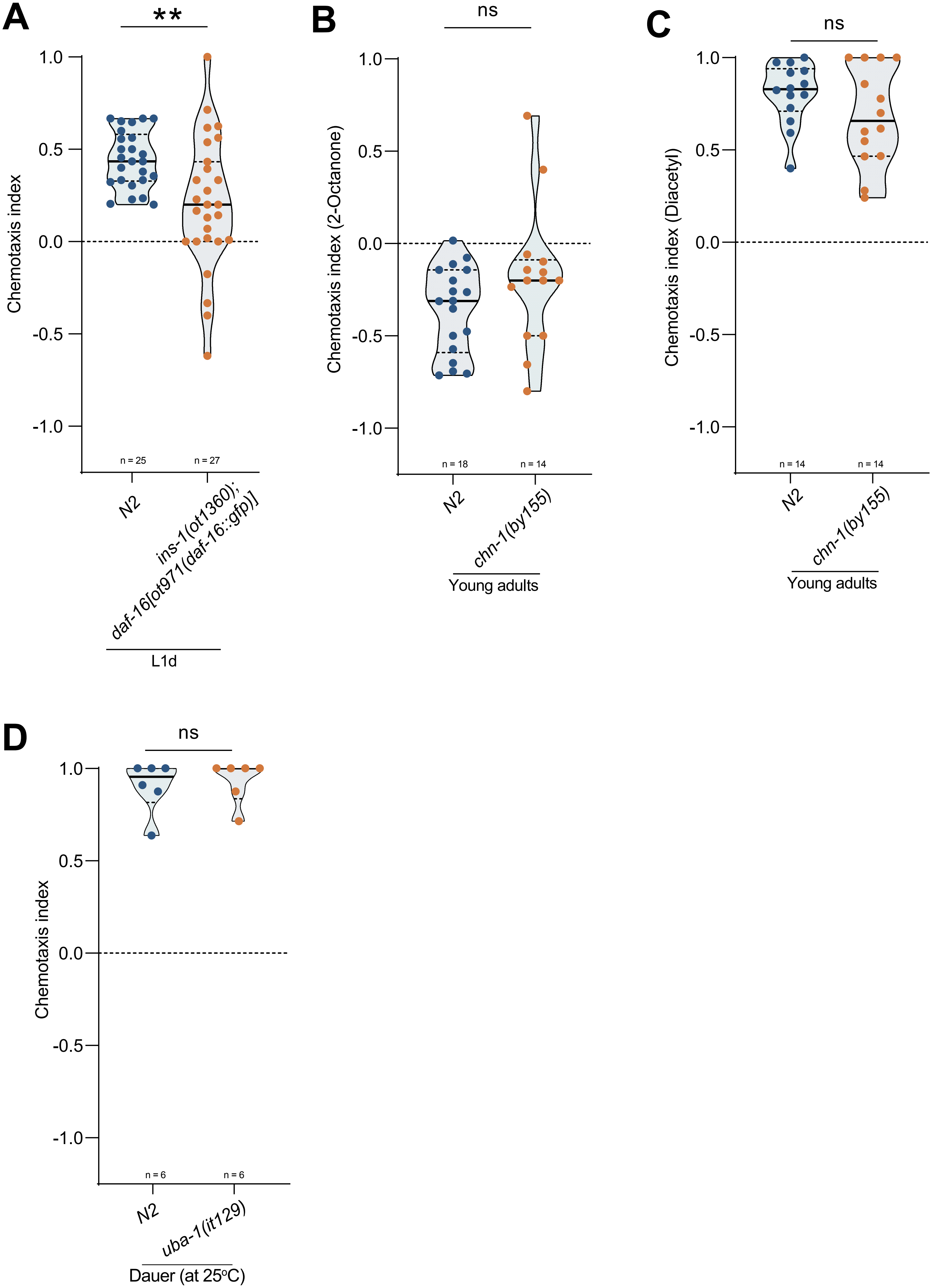

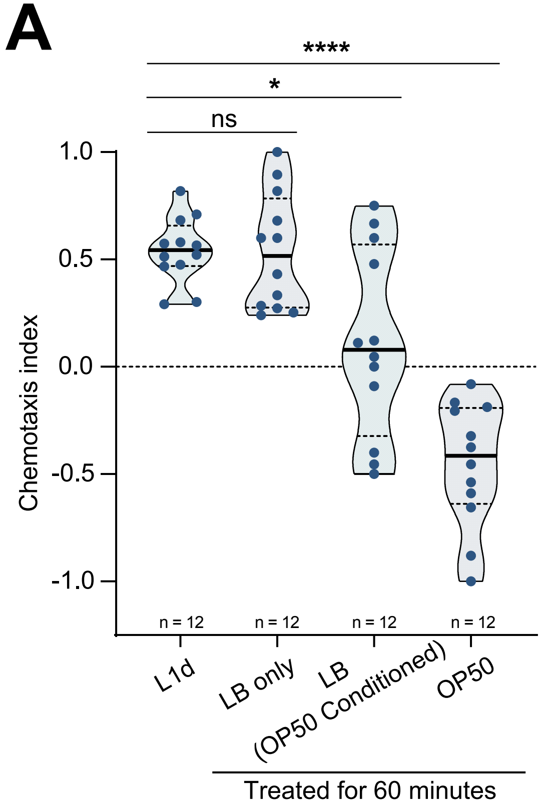

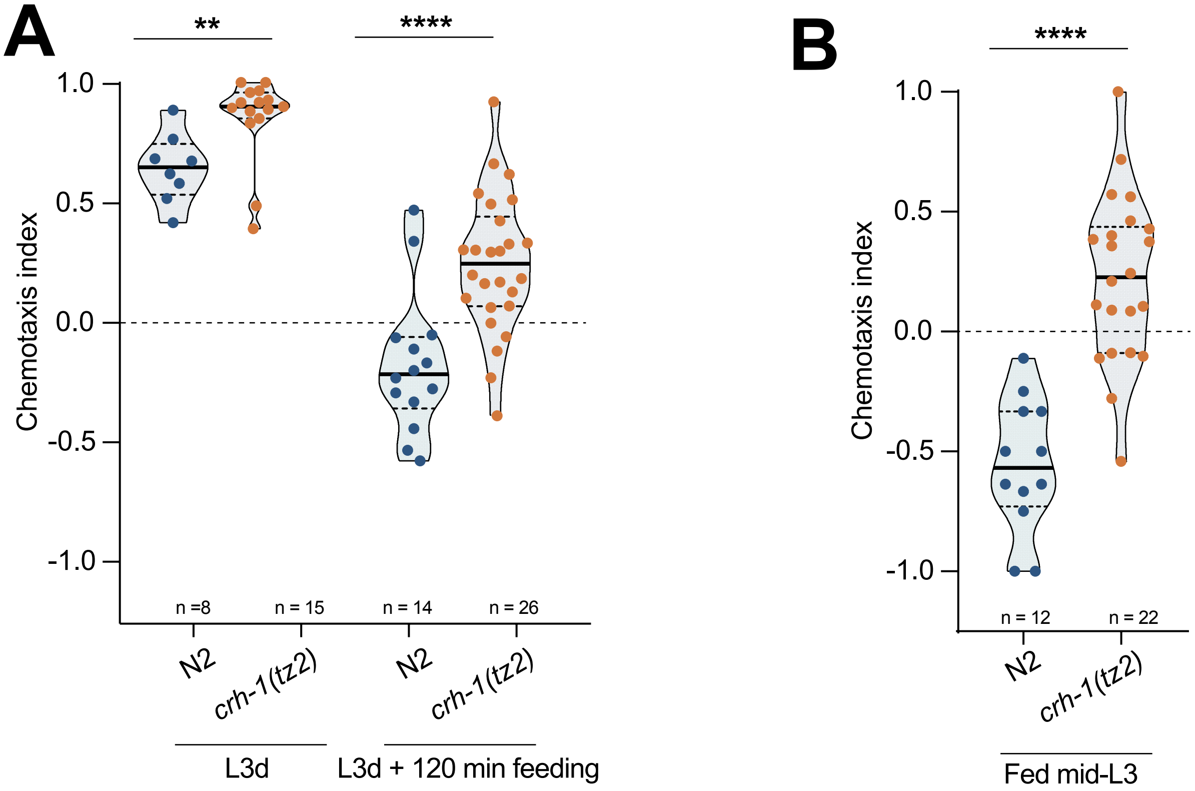
